# The order Tethyida (Porifera) in California: taxonomy, systematics, and the first member of the family Hemiasterellidae in the Eastern Pacific

**DOI:** 10.1101/2020.04.05.026518

**Authors:** Thomas L. Turner

## Abstract

Sponges (phylum Porifera) are ubiquitous inhabitants of marine ecosystems and have been shown to provide substantial ecosystem services. Despite this ecological importance, the sponge fauna in California has received little study. Here I use a collection of kelp forest sponges to describe the diversity of the order Tethyida in California. This order contains the genus *Tethya*, which has been included in long-term ecological monitoring projects. I show that *Tethya vacua* is a junior synonym of *Tethya californiana*, and thus all *Tethya* populations known in California belong to a single species. Genetic data from California’s only Timeidae, *Timea authia*, indicate that this species is in fact in the Tethyidae and is the third known member of the genus *Tethytimea*. I also describe the first member of the family Hemiasterellidae from the Eastern Pacific, *Galaxia gaviotensis* gen. nov. sp. nov. By combining field photographs, morphological taxonomy, and phylogenetic analysis of these samples, this work will facilitate future efforts to understand the evolution of this order and the ecological role of sponges in the California kelp forest.

## Introduction

Though little studied in California, sponges have been shown to play many important roles in other ecosystems, including habitat creation, biological disturbance, and consumption of bacterial and viral plankton (Bell 2008). Sponges in the genus *Tethya* are often called “puffball” sponges, as they grow in a roughly spherical form. Because they are common and distinctive, they are better known than most sponges in California. Indeed, *Tethya* populations – referred to as *T. aurantium* in most ecological publications – have been included in several long-term ecological monitoring efforts aimed at understanding and conserving California’s marine ecosystems (Claisse *et al*. 2018; Miller *et al*. 2018). The Kelp Forest Community Monitoring project, for example, tracks the abundance of 68 diverse marine species at permanent transects in the Channel Islands National Park; *Tethya* are the only sponge included in the survey (Kushner *et al*. 2013). In the 16 sites monitored yearly since the 1980s, *Tethya* were found at every site at least once, and were continuously present in 9 of 16 sites (as of 2011; Kushner *et al*. 2013). In some places they were abundant, with densities up to 0.43 sponges per square meter (Kushner *et al*. 2013). This long-term project also collected presence vs. absence data for over 270 species of animals and algae, and *Tethya* were among the top 10% most frequently present species across all sites (Graham 2004). Though the ecological role of *Tethya* in these habitats remains to be determined, the abundance and ubiquity of these sponges suggests they are ecologically important. Moreover, a large-scale field experiment recently found that *Tethya* populations respond dramatically to kelp forest disturbance. In a spatially replicated field experiment conducted over 9 years, large plots were seasonally cleared of giant kelp (*Macrocystis pyrifera*) to mimic natural disturbance (Castorani *et al*. 2018). The abundances of 205 species of animals and algae were monitored as response variables. *Tethya* biomass increased over 7-fold in plots with increased disturbance frequency, which was the greatest fold-change of all the diverse invertebrates, fish, and algae that were monitored. The reason for this change is unknown, but it motivates additional investigation of this taxon.

Despite this extensive monitoring of *Tethya* populations, it was previously unclear if the monitored populations included one or several species. *Tethya* from Southern and Central California were examined by de Laubenfels (de Laubenfels 1932), who described them all as *T. aurantia var. californiana*, a new variety of the Atlantic species *T. aurantium* (Pallas 1766). The variety was elevated to species status as *T. californiana* in 1993 (Sarà & Corriero 1993). More recently, Austin *et al*. (2014) described a second species of *Tethya* from California, *T. vacua*. This species was described from a single collection at Santa Rosa Island, part of the Channel Islands National Marine Sanctuary in Southern California. This sample was the only one Austin *et al*. examined from Southern California, so they proposed that *T. californiana* is found from British Columbia to Central California, while *T. vacua* is found in Southern California (South of Point Conception, see figure 1). *Tethya* have also been collected from the Gulf of California on Mexico’s West Coast (Sará *et al*. 2001); Austin *et al*. (2014) did not examine these samples but based on published morphological data suggested they may represent an additional yet-to-be described species of *Tethya*, which could also range into California. It was therefore unclear how many species are found in California, whether they can be distinguished in the field, and what their ranges are.

**Figure 1.**
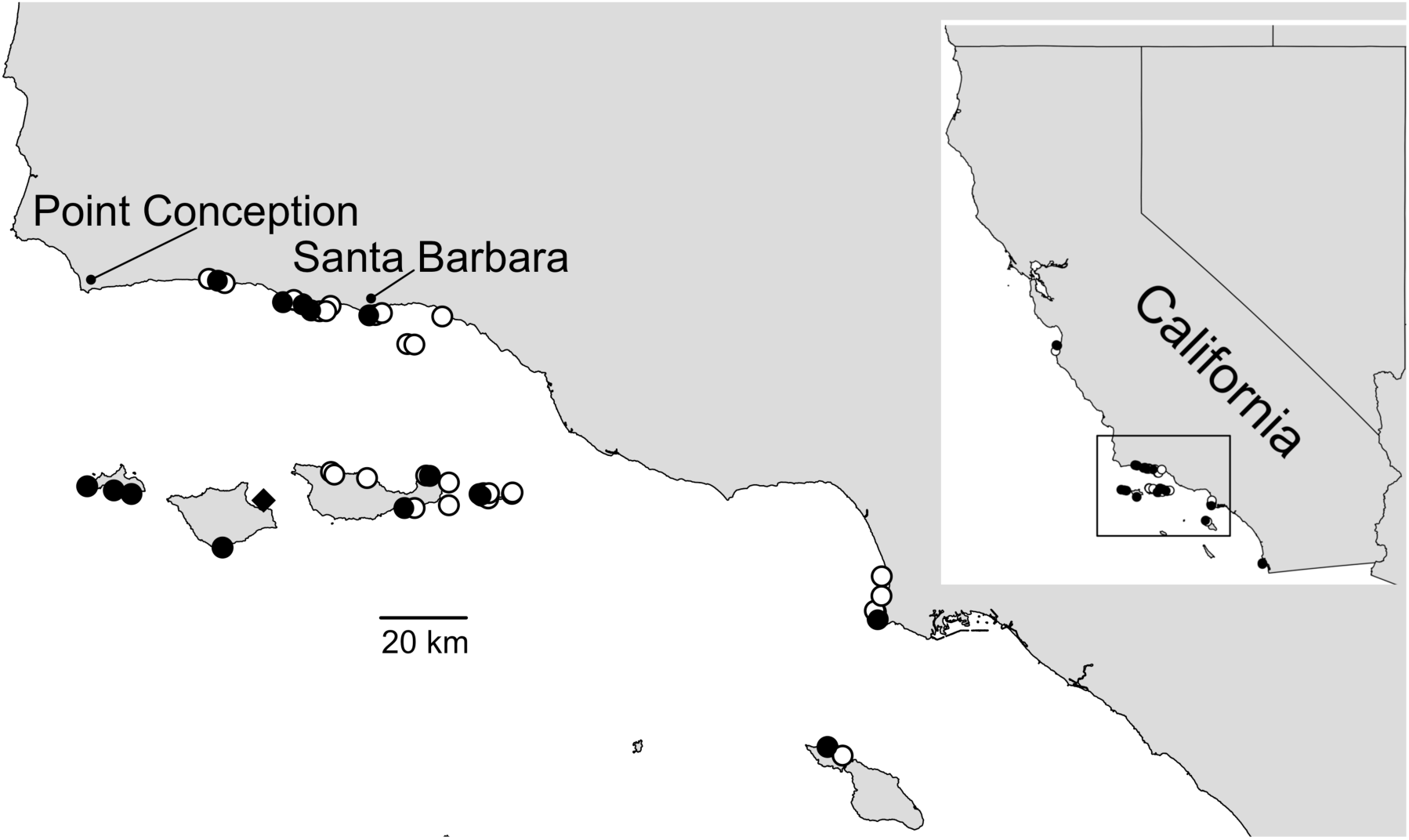
Subtidal collection locations. Sites where *Tethya* were found (black) and not found (white) are shown. The two sites away from the coastline are oil platforms. The diamond indicates the type locality for *T. vacua*.

Here I improve our understanding of these sponges using data from a SCUBA-based survey of kelp forest sponges in California. I have collected and examined the spicules of over 330 sponge samples to date; here I will present the results for all sponges in the order Tethyida. By combining these fresh collections with museum samples and publicly available data, I show that all *Tethya* known in California belong to a single species, *T. californiana*.

The only other sponge from the order Tethyida previously known from California is a member of the family Timeidae: *Timea authia* (de Laubenfels 1932). In contrast to *Tethya*, which are large, conspicuous, and distinctive, *Timea* are encrusting and considerably less apparent. Genetic data from other species of *Timea* have previously revealed this genus to be problematic, with genotyped *Timea* scattered across several different families and even orders (Cruz-Barraza *et al*. 2017, 2020). I provide the first genetic data from *T. authia* and show that it is, in fact, in the family Tethyidae, genus *Tethytimea*.

The third family in the order Tethyida is the Hemiasterellidae. These sponges, nearly all of which have massive, vase-shaped, or upright branching morphologies, are known primarily from the South Pacific and Indian Ocean. Several species are known from Atlantic waters as well, but to my knowledge none have been found in the Eastern Pacific (North or South America). I discovered a new species in Southern California, here formally described as *Galaxia gaviotensis*. Genetic data confirm its membership in the Hemiasterellidae; as a thinly encrusting species, it is morphologically divergent from all previously described species in the family, so I created a new genus to contain it.

## Methods

### Collections

Sponges were located while SCUBA diving. All but two subtidal locations were shallow rocky reefs; the remaining two were oil platforms. Though the survey was focused on kelp forest habitats, I also checked three intertidal sites and floating docks in four harbors, as shown in table S1. I attempted to photograph all sponge morphotypes found at each location, so that presence/absence data could be compared across sites. As shown in table S1, some locations were explored more thoroughly than others. The search times listed are the total dive time, cumulative across all dives at a site. This is only an estimate of search effort, as some dives were spent mainly searching and photographing sponges, while on others considerable time was spent collecting samples. Because of the stochastic nature of the survey, these data should be used to form hypotheses, rather than strong conclusions, regarding species distributions.

Collections were made by hand with a knife. Samples were placed individually in ziplock bags while underwater, accompanied with copious seawater. These bags were put on ice until preservation, which was generally within 2-5 hours, but sometimes up to 12 hours later. Samples were moved directly from seawater to 95% ethanol; in most cases, the preservative was changed with fresh 95% ethanol after 1-3 days, and sometimes changed again if it remained cloudy. Most samples were photographed underwater with an Olympus TG5 before collection and photographed again in the lab. These photos accompany this paper as supplementary data, have been archived with vouchered specimens, and are also posted as georeferenced records on the site iNaturalist.org. Seven samples were vouchered with the California Academy of Sciences in San Francisco; collection numbers are listed in table S2.

### Spicules

To examine spicules, a sponge subsample was placed in a 1.5 ml microcentrofuge tube with household bleach for several hours, until tissue appeared to be dissolved. With the spicules settled at the bottom of the tube, the bleach was then pipetted off and replaced with distilled water; this was repeated several times (I found that 2-3 times was sufficient for visualizing spicules with a light microscope, but preparing samples for scanning electron microscopy required 5 or more rinses and worked best when the final ones were done with absolute ethanol). In some cases, samples were centrifuged at low speed to reduce settling time between rinses, though this increased the proportion of broken spicules.

Spicules were imaged using a compound triocular light microscope and pictures were taken using a D3500 SLR camera (Nikon) with a NDPL-1 microscope adaptor (Amscope). Pictures of a calibration slide were used to determine the number of pixels per mm, and 20-30 spicules were then measured using ImageJ (Schneider *et al*. 2012). For scanning electron microscopy, a sample of spicules in ethanol was pipetted onto a glass microscope slide and allowed to dry. The slide was then coated with ∼20 nm of carbon. Secondary electron images were then taken with a FEI Quanta400F Mk2. Spicule length (or diameter) was measured as the longest possible straight line from tip to tip, even when spicules were curved or bent. The complete distribution of measured values for each sponge is included in the supplementary material.

I also imaged the spicular architecture in cleared tissue sections. I used a razor blade to cut perpendicular sections that were as thin as possible by hand. These sections were prepared in one of two ways. Some were moved from 95% to a 100% ethanol bath, then cleared using Histoclear (National Diagnostics). Other sections were digested in a mixture of 97% Nuclei Lysis Solution (Promega; from the Wizard DNA isolation kit) and 3% 20mg/ml Proteinase K (Promega). Many additional photos of spicules and sections are available in the supplementary data that accompanies this paper, are archived with museum vouchers at the Cal Academy, and have been linked to georeferenced records at iNaturalist.org.

### Genotyping

DNA was extracted using the Wizard DNA extraction kit (Promega), the Qiagen Blood & Tissue kit, or the Qiagen Powersoil kit. All kits performed well on these samples in the sense that downstream PCR was highly successful with all methods. I attempted to amplify fragments of the cox1 mitochondrial barcoding locus for many samples. I used the Folmer primers LCO1490 (GGT CAA CAA ATC ATA AAG AYA TYG G) and HCO2198 (TAA ACT TCA GGG TGA CCA AAR AAY CA) (Folmer *et al*. 1994), which worked for all samples except *T. authia*. For this species, I also tried the primers from Rot et al. (Rot *et al*. 2006), again without success. I also amplified portions of the 28S rDNA nuclear locus. For *T. authia* and *G. gaviotensis*, I amplified the C region using primers C2 (GAA AAG AAC TTT GRA RAG AGA GT) and D2 (TCC GTG TTT CAA GAC GGG) (Chombard *et al*. 1998). These primers amplified bands of multiple sizes for multiple *Tethya* vouchers, but I was able to amplify a larger fragment containing the C region in *Tethya* using primers for the D1-D2 region: Por28S-15F (GCG AGA TCA CCY GCT GAA T) and Por28S-878R (CAC TCC TTG GTC CGT GTT TC) (Morrow *et al*. 2012). For some samples, a portion of the 18S locus was also amplified using the primers SP18aF (CCT GCC AGT AGT CAT ATG CTT) and 600R18S (CGA GCT TTT TAA CTG CAA) (Redmond *et al*. 2013). All primer sequences are listed 5’ to 3’.

PCR was performed in a Biorad T100 thermocycler with the following conditions: 95C for 3 min, followed by 35 cycles of 94C for 30 sec, 52C for 30 sec, 72C for 1 min, followed by 72C for 5 minutes. The 28S C region was amplified with a 57C annealing temperature instead of 52C. PCR was performed in 50 μl reactions using the following recipe: 24 μl nuclease-free water, 10 μl 5x PCR buffer (Gotaq flexi, Promega), 8 ul MgCl, 1 μl 10mM dNTPs (Promega), 2.5 μl of each primer at 10 μM, 0.75 bovine serum albumin (10 mg/ml, final conc 0.15 mg/ml), 0.25 μl Taq (Gotaq flexi, Promega), 1 μl template. ExoSAP-IT (Applied Biosystems) was used to clean PCRs, which were then sequenced by Functional Biosciences using Big Dye V3.1 on ABI 3730xl instruments. All PCR products were sequenced in both directions, and a consensus sequence was constructed using Codon Code v.9 (CodonCode Corporation). Blastn was used to verify that the resulting traces were of sponge origin. All sequences have been deposited in Genbank; accession numbers are listed in table S2.

### Genetic analysis

I used the NCBI taxonomy browser to compile all data from samples identified as belonging to the order Tethyida. I also used blastn to search for additional sequences that appeared to fall within this order. Some accessions identified as Tethyida were then excluded because they were more closely related to other orders of sponges or were from a different region of the gene being analyzed.

Supplementary table S4 lists every Genbank accession found and explains the reasons why any were excluded. Together, included data are from 23 different publications and several unpublished datasets (Chombard 1998; Cruz-Barraza *et al*. 2017, 2020; Dat *et al*. 2018; Erpenbeck *et al*. 2007, 2012, 2016; Heim & Nickel 2010; Heim *et al*. 2007c; a; b; Idan *et al*. 2018; Kober & Nichols 2007; Lavrov *et al*. 2005; Morrow *et al*. 2012; Nichols 2005; Redmond *et al*. 2013; Regueiras *et al*. 2019; Riesgo *et al*. 2013; Shaffer *et al*. 2018; Sorokin *et al*. 2019; Thacker *et al*. 2013; Zhang *et al*. 2016). Sequence alignments were produced in Codon Code v.9 (CodonCode Corporation). Phylogenies were estimated with maximum likelihood using IQ-Tree (Nguyen *et al*. 2015; Trifinopoulos *et al*. 2016). I used the Ultrafast bootstrap (Hoang *et al*. 2018) to measure node confidence. Phylogenies were produced from the IQ-Tree files using the Interactive Tree of Life webserver (Letunic & Bork 2019). Figures were made ready for publication using R (r-project.org) and/or Gimp (gimp.org).

## Results

I used SCUBA to investigate the sponge fauna at 47 sites in Southern and Central California (figure 1, table S1). The spicular morphology of 334 sponges from this project have been analyzed to date, and several were found to belong to the order Tethyida. After follow-up collections targeting specific species and locations, a total of 13 specimens could be assigned to the Tethyida. Below, I explain the results of the phylogentic analysis of these samples, and what these phylogenies tell us about other taxa. This is followed by a systematic section with additional details on each species.

### Genetic Results

To perform genetic analysis of newly collected samples, I focused on the three loci most frequently used in sponge systematics: cox1 and the large (28S) and small (18S) ribosomal subunits. As detailed in the methods, not all loci were successfully amplified from all species. When possible, I sequenced sponges from both Central and Southern California for comparison. I then compiled all published records in Genbank for the order Tethyida for comparative analysis (see methods).

Of the three loci investigated, the largest number of public records were available at the 28S rDNA (figure 2). Considering only the previously published data, the phylogeny is very similar to a recently published phylogeny (Sorokin *et al*. 2019), though my interpretation of these results differ on several key points. In addition, I excluded records that lacked the highly variable C region. Genbank records at 28S differ greatly in read length: though phylogenetic methods can accommodate missing data, it is preferable to have at least a single region shared by all records. Though this led to the exclusion of several species, it also improved the stability of the phylogeny. One notable improvement is the placement of *T. californiana*: the published phylogeny of Sorokin et al. (2019) found that *T. californiana* nested within the European *T. hibernica/citrina/norvegica* clade at 28S, but was in the sister group to this clade at cox1. The 28S sequence used in their analysis did not include the C2 region. I sequenced an 824 bp region that did include C2 from a freshly collected *T. californiana* from Southern California, and this sequence has a phylogenetic position consistent with what is found in other gene trees.

**Figure 2.**
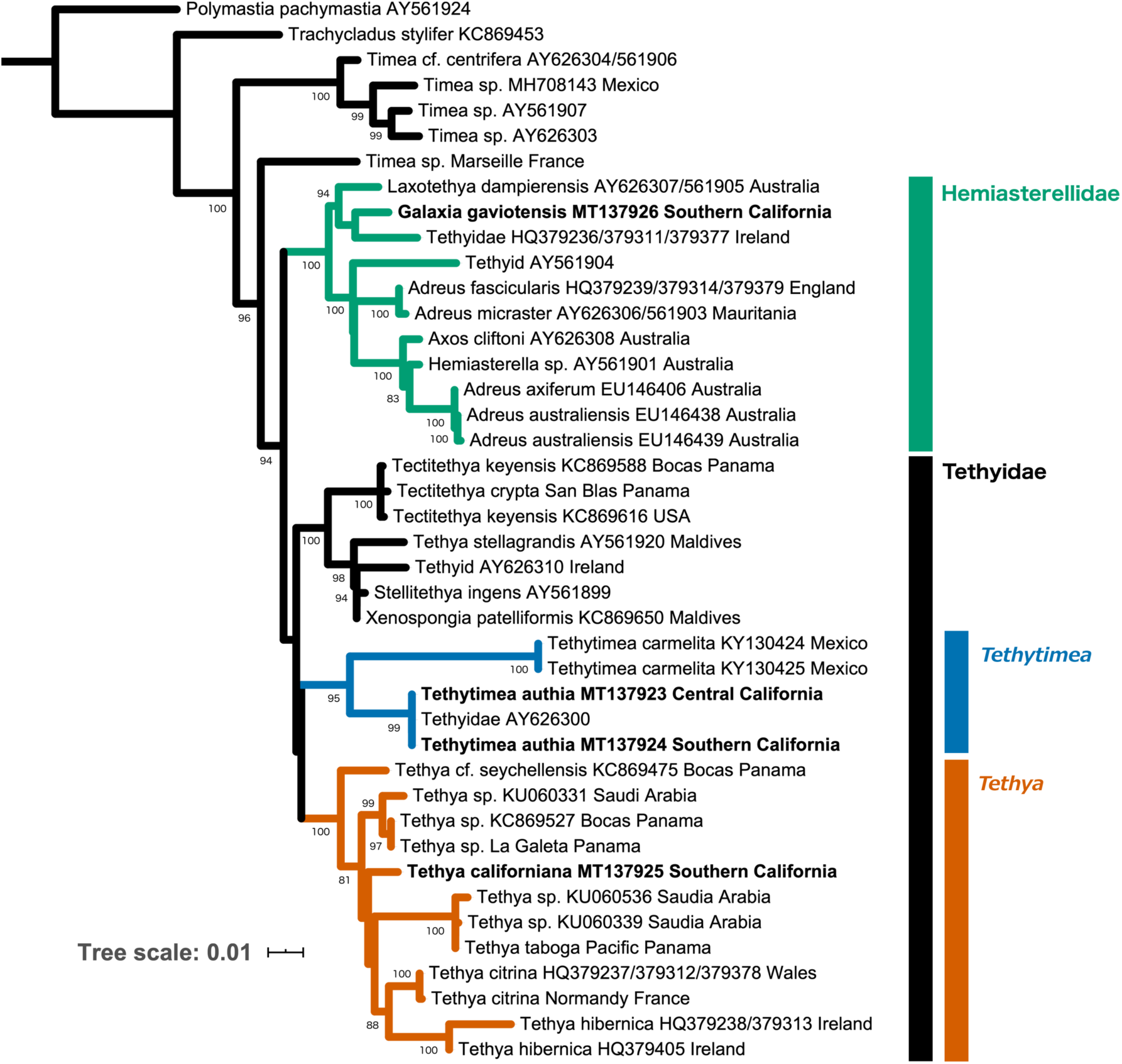
Gene tree at the large ribosomal subunit (28S). Bootstrap values are shown for nodes with > 80% support; nodes with < 50% support are collapsed. Green = F. Hemiasterellidae, blue = *Tethytimea*, orange = *Tethya*. Genbank accession numbers are shown; those beginning with MT, shown in bold, are new. Scale bar indicates substitutions per site.

The 28S phylogeny recovers the Family Tethyidae as monophyletic, if the systematic placement of two taxa are revised (as detailed below). Three well supported clades of Tethyidae are recovered. One clade includes all records from the genus *Tethya* save one: *T. stellagrandis*. This species falls into a second well supported clade that includes nearly all other Tethyidae with available data (eight additional genera have no available sequence data at this locus). The third and final clade consists of *Tethytimea carmelita* and an unidentified sample. The unidentified sample is from a voucher that is now lost, but personal communications with the collector, Scott Nichols, indicate it was likely from Central California. I sequenced the 28S locus in two newly collected individuals of *Timea authia*: one from Central California (Monterey Bay) and one from Southern California (Carpinteria Reef). Both of these sequences are identical to the unidentified sample over the aligned 816 bp, so it is likely the unidentified sample is also of this species. These three samples form a clade with *Tethytimea carmelita* with 95% bootstrap support. This result, together with the genetic data from 18S and the morphological data below, leads me to reassign *Timea authia* to the genus *Tethytimea* (as detailed in the Systematics section).

Only one species of Tethyidae is not included in the 28S clade discussed above: *Laxotethya dampierensis*, which is the type species of its genus. This species falls into the sister clade to the Tethyidae. This sister clade is itself composed of two clades: one that includes all species of the Hemiasterellidae with available data, and another that includes *L. dampierensis*, an undescribed species identified as “Tethyidae” in Genbank, and *Galaxia gaviotensis* gen. nov. sp. nov.. The “Tethyidae” sample, collected in Ireland, is in the process of being described by others. Based on a personal communication with Bernard Picton, the collector, it is encrusting and has some traits in common with *Adreus* (F. Hemiasterellidae), but does not fit within any known genus. Based on the phylogenetic position of this clade, the simplest way to resolve its higher taxonomy would be to move *Laxotethya* to the Hemiasterellidae, making this family the sister clade to the Tethyidae (figure 1). I formally suggest this reassignment here (though I did not examine the *L. dampierensis* voucher that was sequenced, I did verify that it was likely to be correctly identified; it is present at the Western Australia Museum as voucher WAMZ11871, identified by Dr. Jane Fromont). This would place the undescribed Irish sponge and the new California sponge in the family Hemiasterellidae as well.

The reassignment of *Timea authia* adds to a growing list of taxonomic problems for the genus *Timea* (Cruz-Barraza *et al*. 2017, 2020), which is currently in its own family (Timeidae). The only other *Timea* record at 28S that is identified to species, and includes the C region, is *Timea lowchoyi* (AY561871). This sample is not shown on the phylogeny, because it is nested within the order Axinellida (Sorokin *et al*. 2019). The remaining samples, all *Timea sp*., do not form a single clade. These results could indicate that 1) samples have been misidentified, or 2) the family is highly polyphyletic and in need of revision. My results for *T. authia*, and the published results of others, suggest that the need for revision is real, even if some samples have been misidentified (Cruz-Barraza *et al*. 2020).

The gene tree of the Folmer barcoding region of the cox1 gene is consistent with the key results of the 28S tree (figure 3). The family Hemiasterellidae forms a single clade that includes *G. gaviotensis* sp. nov. The genus *Tethya* again forms a well supported clade; unfortunately, no sequences from Tethyidae outside of Tethya are available thus far (attempts to amplify this locus from *T. authia* repeatedly failed). *Timea* sequences from Genbank again illustrate that this genus requires revision. Two sequences are not shown: *Timea unistellata* from Ireland (KC869427) and an unidentified *Timea sp*. (AY561968). Both of these sequences fell within the order Poecilosclerida. This was recently reported by Cruz-Barraza et al. (2020), who show that these two *Timea* are not closely related to each other within that order (their figure 6). The remaining *Timea* sequences do form a well supported clade, but the placement of this clade is different than for the 28S locus. None of the *Timea* vouchers sequenced at cox1 have also been sequenced at 28S, so the most likely explanation is that there are some *Timea* that are outside of the Hemiasterellidae + Tethyidae (as shown in the 28S tree) and other *Timea* that are more closely related to Tethyidae than Hemiasterellidae (as shown in the cox1 tree). Note that the placement of the *Timea* clade in the cox1 phylogeny is consistent with the placement of the morphologically similar genus *Tethytimea* in the 28S phylogeny.

**Figure 3.**
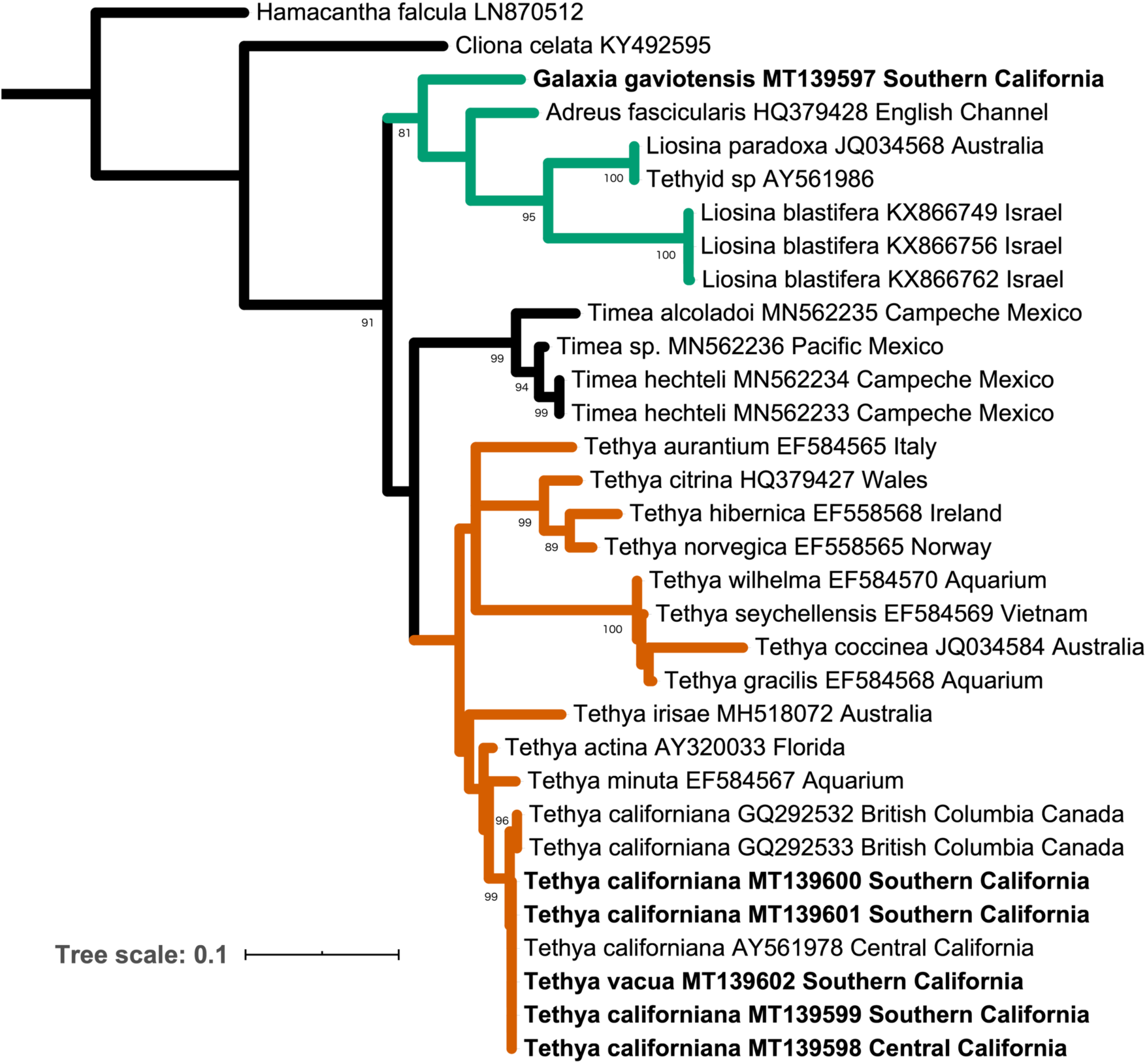
Gene tree at cox1. Bootstrap values are shown for nodes with > 80% support; nodes with < 50% support are collapsed. Green = F. Hemiasterellidae, orange = *Tethya*. Genbank accession numbers are shown; those beginning with MT, shown in bold, are new. Scale bar indicates substitutions per site.

**Figure 4.**
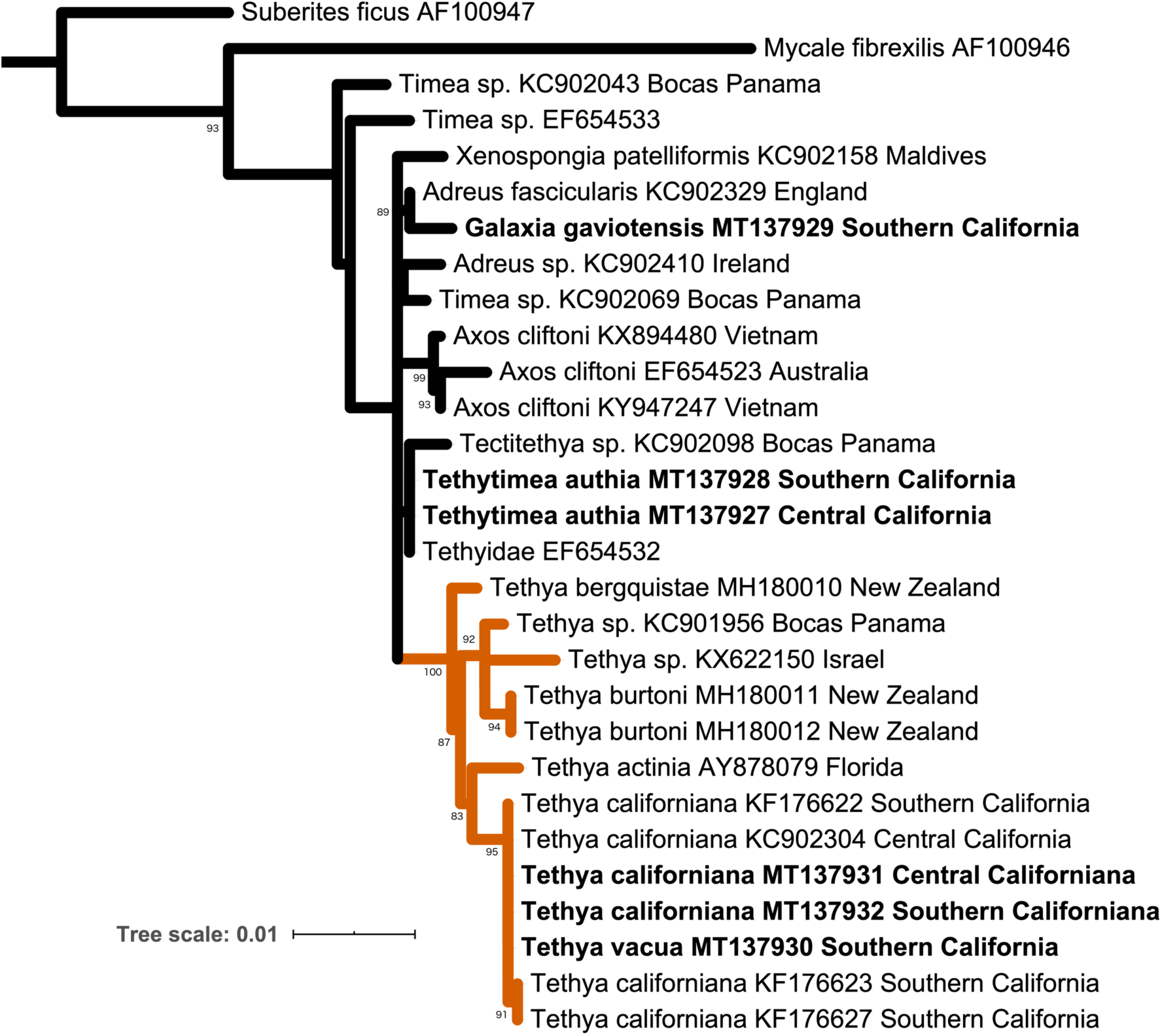
Gene tree at the small ribosomal subunit (18S). Bootstrap values are shown for nodes with > 80% support; nodes with < 50% support are collapsed. Orange = *Tethya*. Genbank accession numbers are shown; those beginning with MT, shown in bold, are new. Scale bar indicates substitutions per site.

**Figure 5.**
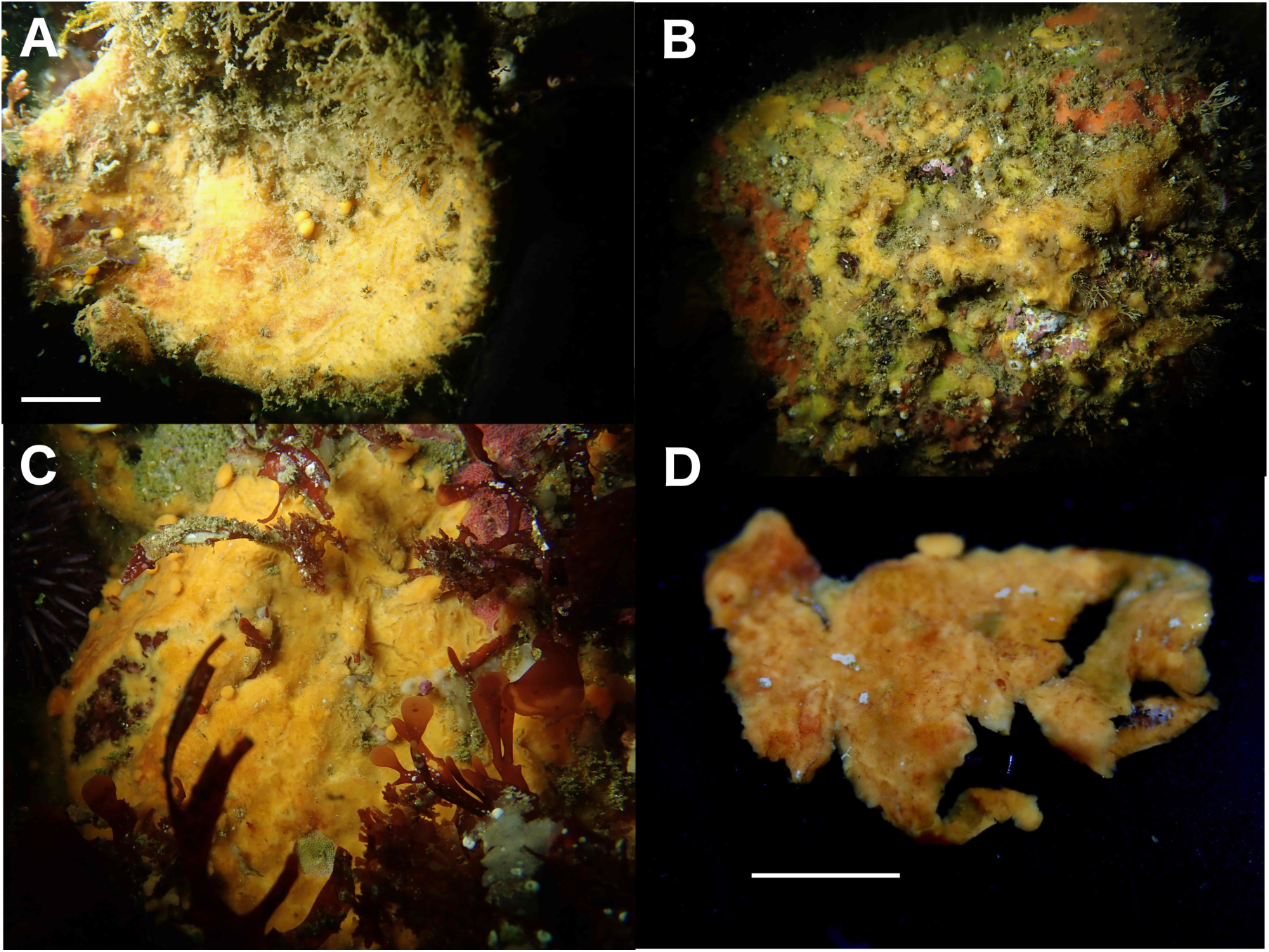
*Tethyatimea authia*. A) CASIZ 235110, Carpinteria Reef; B) TLT 364, Saddleback Ridge, Santa Cruz Island; C) CASIZ 235111, Perkin’s Reef, Monterey Bay; D) CASIZ 235111 after collection but before preservation, showing constriction of gutters. Scale bar = 1 cm.

**Figure 6.**
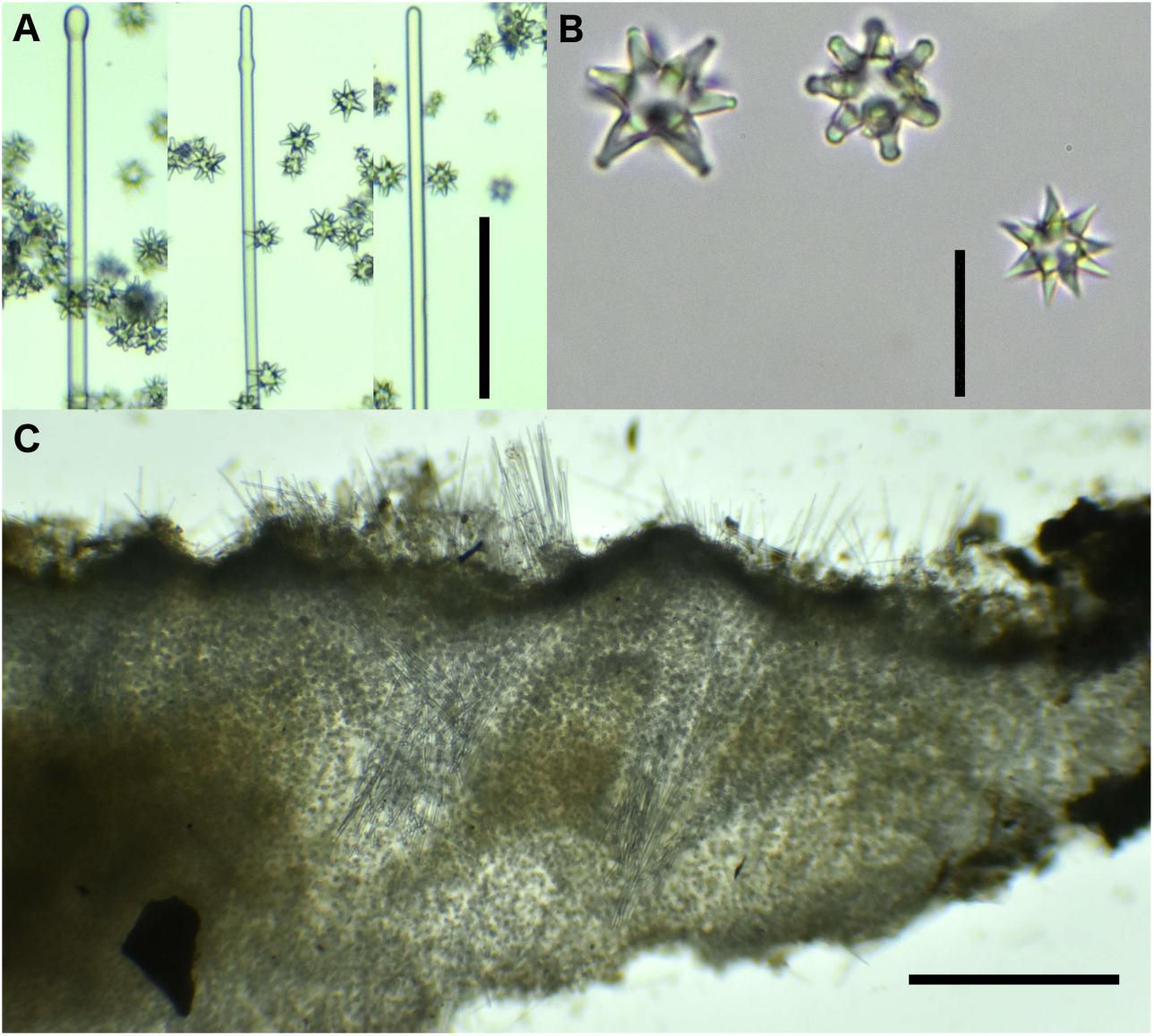
Microscopic features of *T. authia*. A) tylostyles, styles from voucher CASIZ 235111, scale bar 100 μm; B) euasters from voucher CASIZ 235111, scale bar 20 μm; C) perpendicular tissue section from voucher TLT 364, scale bar = 500 μm.

The cox1 phylogeny fails to support species status for *Tethya vacua*. Three sequences were previously available for *Tethya californiana*; two of these are from British Columbia, while the other is from Central California. The Central California sample is somewhat uncertain: it is thought to be from Carmel Bay, California, which contains the type locality for *T. californiana*. However, the voucher and collection information are lost. I therefore collected a new sponge from Monterey Bay, about 8 km North of the type location, and sequenced it for comparison (MT139598 in figure 3). A *T. vacua* paratype was also sequenced, and is genetically identical to the Monterey sponge at the 650 bp Folmer region of cox1. Three additional sponges from island and mainland locations in Southern California were likewise identical. The British Columbia sponges are slightly divergent from this California haplotype (0.6% absolute sequence divergence). This divergence was previously noted vs. the Central California sponge from Carmel, and was used to support species status for this population under the name *T. leysae* (Heim & Nickel 2010). Austin *et al*. (2014) synonymized *T. leysae* with *T. californiana;* they argued that this level of genetic divergence provides limited support for species status.

The phylogeny at the 18S locus is less resolved than the other loci due to the slow evolutionary rate at this locus (figure 4). I sequenced ∼850 bp from *Tethya californiana* from Central and Southern California, and again found that they had identical haplotypes. I was able to obtain a homologous 600 bp from the *T. vacua* paratype and it was identical as well. Furthermore, a 1600 bp region was previously sequenced from several *Tethya* from Naples Reef (Southern California) and the *T. californiana* that is likely from Carmel (Central California); one of these Southern California sponges is identical to the Carmel sponge over this entire region, while the others are nearly identical. These data again fail to support species status for *T. vacua*, nor do they support genetic differentiation between Southern and Central California in this genus.

The 18S phylogeny is consistent with the placement of *T. authia* in the Tethyidae. No other *Tethytimea* have been sequenced at this locus, but *T. authia* sequences form a (poorly supported) clade with the only Tethyidae sequences available outside of the genus *Tethya*. Note that that there is a published sequence that is identical to *T. authia* at 18S (EF654532): this is from the same voucher that is identical at 28S, which is likely to have been a *T. authia*.

The 18S phylogeny also supports the placement of *G. gaviotensis* sp. nov. in the Hemiasterellidae. The most supported tree at 18S placed all Hemiasterellidae in a single clade, but this clade had only 33% bootstrap support, and is collapsed in figure 4. Despite this, *G. gaviotensis* sp. nov. clusters with *Adreus fascicularis* with good support, consistent with the 28S phylogeny.

The 18S phylogeny once again illustrates the need for revision of *Timea*. Though all three available sequences grouped within the Tethyida, two are outside the Hemiasterellidae + Tethyidae, while the other is weakly supported as within the Hemiasterellidae.

Finally, these phylogenies indicate something potentially interesting regarding *Axos cliftoni*. This species is placed within the Hemiasterellidae, and sequences at 18S and 28S are consistent with this placement. Sequences at cox1, however, are not shown, because preliminary analyses showed them to group in the order Axinellida. This was found by Cruz-Barraza et al. (2020) as well, as shown in figure 6 of their paper. This is perplexing, because the cox1 sequences are annotated as being from the same vouchers as the rDNA sequences, and come from different collections in two countries. Voucher MT5.2015 is from Vietnam and was sequenced at two loci (18S = KY947247, cox1 = KY947263). Voucher G300111 is from Australia and was sequenced at three loci (28S = AY626308, 18S = EF654523, cox1 = AY561974). The cox1 sequences from these two samples are extremely similar, but group with family Stelligeridae in the order Axinellida (Cruz-Barraza *et al*. 2020). The 18S sequences from these vouchers are similar, and form a clade within the Tethyida (figure 4); the 28S sequence from Australia falls within the Hemiasterellidae (figure 3). Perhaps this conflict is due to errors with sequence annotations in Genbank, but alternatively, it may suggest horizontal gene transfer or some other biologically interesting phenomena.

## Systematics

Order TETHYIDA Morrow & Cárdenas, 2015

Family Tethyidae Gray, 1848

Genus *Tethya* Lamarck, 1815

*Tethya californiana* de Laubenfels, 1932

## Synonymy

*Tethya aurantia var. californiana* de Laubenfels, 1932

*Tethya vacua* Austin et al. 2014

## Material Examined

CASIZ 235109, Hopkin’s Marine Station, Monterey CA (34.47182, -120.14262), 9/16/19. CASIZ 235108, Naples Reef, Santa Barbara CA (34.42212, - 119.95154), 7/31/19. CASIZ 235106, Elwood Reef, Santa Barbara CA (34.41775, - 119.90150) 4/17/19. CASIZ 235107, Santa Cruz Island, CA (33.98378, -119.63910), 4/26/19. TLT 361, Santa Rosa Island, CA (33.89966, -120.10735) 10/9/19. CASIZ 184729, Santa Rosa Island, CA (34.00000, -120.00170), collected by France & Efford, 12/24/81 (*T. vacua* paratype). CNPGG 1199, Ensenada, Bahia Topolobampo, Baja California, Mexico, 4/8/11. CNPGG 0012, Cabo San Miguel, Gulf of California, Mexico, 3/5/85. (For tabular format, see supplemental table 2).

**Table 2.** Spicule data for *Tethytimea authia* in microns.

|  | Carpinteria Reef, Santa Barbara CA | Perkin's Reef, Monterey CA | Saddleback Ridge, Santa Cruz Island, CA | Timea authia (Carballo & Cruz-Barraza 2006) |
| --- | --- | --- | --- | --- |
| Survey ID | TLT 125 | TLT 449 | TLT 364 | - |
| Museum Voucher | CASIZ 235110 | CASIZ 235111 | - | - |
| Collection date | 7/31/19 | 11/24/19 | 10/18/19 | - |
| All styles: |  |  |  |  |
| Sample size | 41 | 28 | 35 | - |
| Length | 110-(491)-779 | 309-(560)-812 | 176-(469)-883 | - |
| Width | 2-(5.3)-10 | 5-(7.1)-10 | 3-(5.3)-10 | - |
| Tylostyles only: |  |  |  |  |
| Sample size | 29 | 11 | 19 | - |
| Length | 208-(543)-779 | 379-(653)-797 | 176-(475)-748 | 208-(478)-915 |
| Width | 2-(5.3)-9 | 5-(7.8)-10 | 3-(5.4)-9 | 4-(6.4)-15 |
| Simple styles only: |  |  |  |  |
| Sample size | 11 | 12 | 16 | - |
| Length | 110-(372)-555 | 309-(497)-812 | 176-(461)-883 | 187-(289)-800 |
| Width | 2-(5.5)-10 | 5-(6.8)-9 | 3-(5.3)-10 | 3-(3.5)-5 |
| Asters: |  |  |  |  |
| Sample size | 219 | 369 | 204 | - |
| Diameter | 6-(14.8)-25 | 7-(17.7)-26 | 3-(15.5)-28 | 3-(14.7)-24 |

## Description

### Macroscopic features

Approximately hemispherical, with broad gutters between tubercles when relaxed. The sponge constricts upon collection, closing the gutters and placing the tubercles in contact. The width and depth of the gutters varies greatly in field photos, as does color (from yellow to orange, but sometimes covered in a dark red growth of apparent algae). As this species has been well described elsewhere (Austin *et al*. 2014), I focused on improving our understanding of the quantitative aspects of spicules that have been shown to vary among samples.

### Spicules

To assess the possibility of multiple species of *Tethya* within the study region, I quantified two aspects of spicular morphology found to vary in previous studies (Austin *et al*. 2014; Heim & Nickel 2010; Sará *et al*. 2001). The first concerns the morphology of the megascleres. Austin et al. (2014) divided megascleres into three categories defined as follows:

-anisostrongyles: distal end (foot) smaller diameter than proximal end (head).
-strongyloxeas: fusiform, distal end (foot) a point; sides narrow toward rounded end (head).
-styles: straight sides, distal end (foot) a point; diameter of rounded end (head) equal to that of shaft.

After examining a large number of spicules, I found strongyloxeas and styles to be part of a continuous spectrum. I measured the width of each spicule at the head and the widest point, and some were very fusiform (widest point more than 4 times wider than head), some were clearly styles (widest point equal to head), but most spicules were intermediate. I therefore grouped these two types together. Anisostrongyles were easier to distinguish, as many spicules clearly lacked a sharp point, though once again there were spicules that were intermediate and hard to classify. I therefore present data on all megascleres together, and also present data on anisostrongyles separately (table 1). Regardless of method, the distributions of megasclere dimensions broadly overlapped among samples. ANOVA of all megascleres considered together revealed no significant differences among samples (F=1.73, p=0.13). One sample, from the Gulf of California (Mexico), was found to lack anisostrongyles almost completely, while anisostrongyles comprised 27-45% of megascleres in all other samples (see remarks).

**Table 1.**
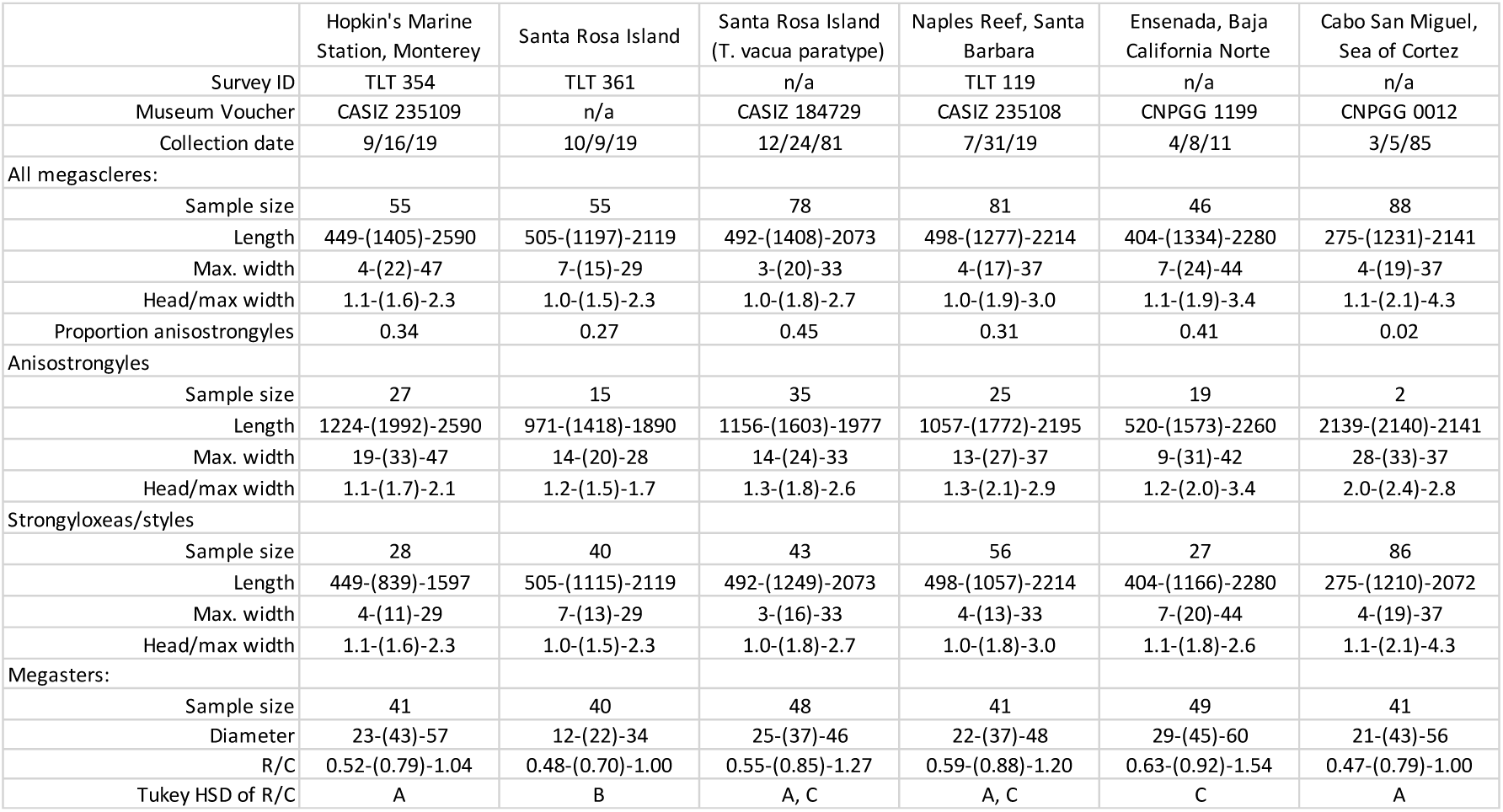
Spicule data for *Tethya californiana* in microns.

The other quantitative measure that has been found to vary among *Tethya* in this region relates to megasters (star-shaped spicules). By dividing the ray length by the centrum diameter, a parameter called R/C can be calculated for each aster (Austin *et al*. 2014). I did this for 40-50 asters per sponge, and used ANOVA to determine if there were significant differences among samples (there were, F=12.14, p=1.81e-10). Tukey’s honest significant differences test was then use to determine which samples were different from others at the 95% family-wide confidence level. Note that the *T. californiana* sample from Hopkin’s Marine Station in Central California (near the type location for *T. californiana*) was not significantly different from the *T. vacua* paratype (table 1).

## Remarks

Though *Tethya* are the best studied sponge in California from an ecological perspective, their taxonomy has remained unresolved. *Tethya aurantia* var. *californiana* de Laubenfels was described in 1932, with Carmel Bay, in Central California, as the type location (de Laubenfels 1930). The variety was raised to species status in 1993 (Sarà & Corriero 1993). Heim and Nickel (2010) then split this taxon, describing those from British Columbia as a separate species, *T. leysae. Tethya leysae* was synonymized with *T. californiana* by Austin *et al*. (2014), who proposed that this species ranged from British Columbia to Central California. In the same monograph, Austin et al. erected *T. vacua* for Southern California, based on a single collection from Santa Rosa Island (figure 1). They also implied that the sponges described as *T. californiana* from the Gulf of California, at the Southern range limit, were likely a different species. I wished to confirm or refute the validity of the name *T. vacua*, and assess the possibility of an additional species in the South.

Genetic data do not support species status for *T. vacua*. In the same monograph where *T. vacua* was described, Austin et al. (2014) argued that 0.6% absolute sequence divergence at cox1 was insufficient to support species status for *T. leysae*. I show that *T. vacua* are even more similar (0% sequence divergence) to *T. californiana* at this same locus. The same is true at the 18S nuclear locus.

The spicules of Central and Southern California *Tethya* are likewise indistinguishable. The only spicule data used to support *T. vacua* was that the one sample analyzed had asters with a higher R/C value than the *T. californiana* syntype. No statistical test was used, however, and another *T. californiana* sample from Monterey Bay (CASIZ 067731, table 14 of Austin *et al*. 2014) was very similar to the *T. vacua* sample. Here I show that the R/C values of the *T. vacua* paratype are not significantly different from another sponge collected in Monterey Bay, near the type location for *T. californiana*. Though not statistically significant, average values are slightly larger for 2 of the 3 Southern California samples analyzed here vs. the Central California sample, and this is consistent with a geographic cline in R/C value as previously reported (Austin *et al*. 2014). Austin et al. show that R/C values are shortest in cold British Columbian waters, and increase towards the South (and again used this cline to argue against species status for *T. leysae*). R/C values of the Ensenada sponge analyzed here have the largest values, consistent with this cline.

The only other morphological difference reported for *T. vacua* are the vacuoles that are its namesake. I compared tissue sections of sponges from Monterey (Central California), Naples Reef (Southern California), and Santa Rosa Island (Southern California, near the holotype location for *T. vacua*), to look for these vacuoles (see supplementary data). I failed to find vacuoles in any sponge, but these negative data should not be considered conclusive. I did not have access to the *T. vacua* holotype as a positive control, so it is possible that my methods were inadequate to see them. In any case, I feel that the lack of any spicular characters, combined with the lack of genetic differentiation, clearly indicates that these taxa should be synonymized.

I was also interested in whether there is an additional species of *Tethya* at the Southern end of the *T. californiana* range. Two sponges from Pacific Mexico were graciously shared with me by Patricia Gómez, Porifera Curator at the National Autonomous University of Mexico. One is from Ensenada, just South of the US/Mexico border, where habitats are similar to California kelp forests. The spicules of this sponge were indistinguishable from those in Southern California (table 1). The other was from Cabo San Miguel, in the Gulf of California; this is beyond the range of the giant kelp *Macrocystis pyrifera*, in an ecosystem quite different from those on California shores. Sponges from this collection were described as having several morphological differences compared to the *T. californiana* holotype, the most significant of which was having only strongyloxeas instead of anisostrongyles and strongyloxeas (Sará *et al*. 2001). Sará, Gómez, and Sára (2001) did not feel these differences warranted species status. In my reanalysis of one of these sponges, I replicate their finding of (nearly) exclusive strongyloxeas, and find that the R/C values of asters are somewhat anomalous compared to the clinal expectation. I attempted to sequence fragments of cox1 and 18S from these Mexican samples, but was unable to amplify any DNA (I also attempted to sequence DNA from other archived samples of Mexican *Tethya* species, again without success). Collection of fresh material for sequencing is therefore highly recommended. It remains unconfirmed if the Gulf of California *Tethya* are *T. californiana*, but the similarities of the Ensenada sponge to California samples makes it unlikely that this Gulf of California morphotype range into the state of California.

## Geographic distribution

*Tethya californiana* was previously described as ranging from British Columbia in the North to Central California in the South (Austin *et al*. 2014). Genetic data now extend this range to Southern California, and morphological comparisons extend it to at least Ensenada, Mexico. Consistent with published reports, this sponge was common in kelp forest habitats in California. I observed it at 19/47 (40%) of kelp forest locations surveyed. There are some indications it may be more abundant in colder waters (because of the nature of the survey, these patterns should be considered hypotheses only; see methods). Water temperatures are colder in Central California than Southern California, and within the Santa Barbara Channel there is a strong gradient, with colder waters in the West (Claisse *et al*. 2018). *Tethya californiana* were present at 2 of 3 sites in Central California, and 4 of 4 sites on the Western-most islands of San Miguel and Santa Rosa. In contrast, it was seen at only 2 of 8 location on Anacapa Island, and 2 of 7 locations in Los Angeles County. It was not present at the two oil platforms surveyed, nor on floating docks in the four harbors surveyed. I did not find it in the 3 intertidal locations checked, but pictures posted by others to the site iNaturalist show that the sponge is sometimes present in the intertidal in Northern and Central California (and the holotype collected in Central California in 1926 was an intertidal sponge).

Genus *Tethytimea* de Laubenfels, 1936

*Diagnosis*. Tethyidae with thinly encrusting body. Megasclere skeleton composed of parallel bundles of tylostyles, with or without styles, ending in tubercles. Megasters are oxyspherasters and tylasters, heterogeneous in size and sometimes with giant spicules. Micrasters are tylasters. Other microscleres, occasionally present, are spheres. (Modified from Cruz-Barraza *et al*. (2017) to include styles and tylasters among megasters.)

*Tethytimea authia* de Laubenfels, 1930 (comb. nov.)

## Synonymy

*Timea authia* de Laubenfels, 1930

## Material Examined

CASIZ 235111, Perkin’s Reef, Monterey CA (36.62920, - 121.92031), 11/24/19. CASIZ 235110, Carpinteria Reef, Santa Barbara CA (34.39163, - 119.54169), 7/31/19. TLT 364, Saddleback Ridge, Santa Cruz Island CA (34.03817, - 119.52470), 10/18/19.

## Description

### Macroscopic features

An encrusting sponge; samples vary from 0.5 - 2.0 mm thick. Color yellow/orange when alive (figure 5). The surface is sculpted by gutters in situ, but the sponge contracts when collected, and gutters are no longer visible. Scattered, mushroom-like papillae present on surface, approximately 1-3 mm in diameter. No other sponges were located with these features in the current survey: it is therefore possible that this species can be identified from field photos, at least within the Southern California portion of its range. Note that these characters are difficult to see in the field, and more apparent in macro photos.

### Spicules

Megascleres are tylostyles with oval heads, styles with an enlarged band near the head, and simple, unadorned styles (figure 6). Microscleres are asters. The most common aster has thick, non-tapering rays with enlarged tips, usually with spines at the tips (tylasters). Others have cone-shaped rays ending in sharp tips (oxyspherasters). Intermediates are also found, with tapering rays that are fairly cone-shaped, but ending in enlarged tips with spines. Spicule measurements are shown in table 2, and compared to published data (Carballo and Cruz-Barraza 2006). Carballo and Cruz-Barraza (2006) reanalyzed the holotype and used SEM to provide more detail regarding spicule shape. Data from my three samples are entirely consistent with their data, qualitatively and quantitatively.

*Tethytimea* are described as having two size classes of asters: larger “megasters” and smaller “micrasters”. *Timea* are described as having only “asters”. I quantified the size distribution of asters to determine whether there was more than one size class (figure 7). One sample (TLT364) seemed to have two size classes, with “megasters” 13-30 μm and “micrasters” 3-7 μm (though 2% of asters were intermediate). The other two sponges had two modes, but the distributions of the size classes were overlapping. I used Hardigan’s Diptest to test for unimodality, which was rejected for all three samples (p<0.0001 for all). The larger sized asters included both oxyspherasters and tylasters, with tylasters more common. Only tylasters were seen among the smallest asters.

**Figure 7.**
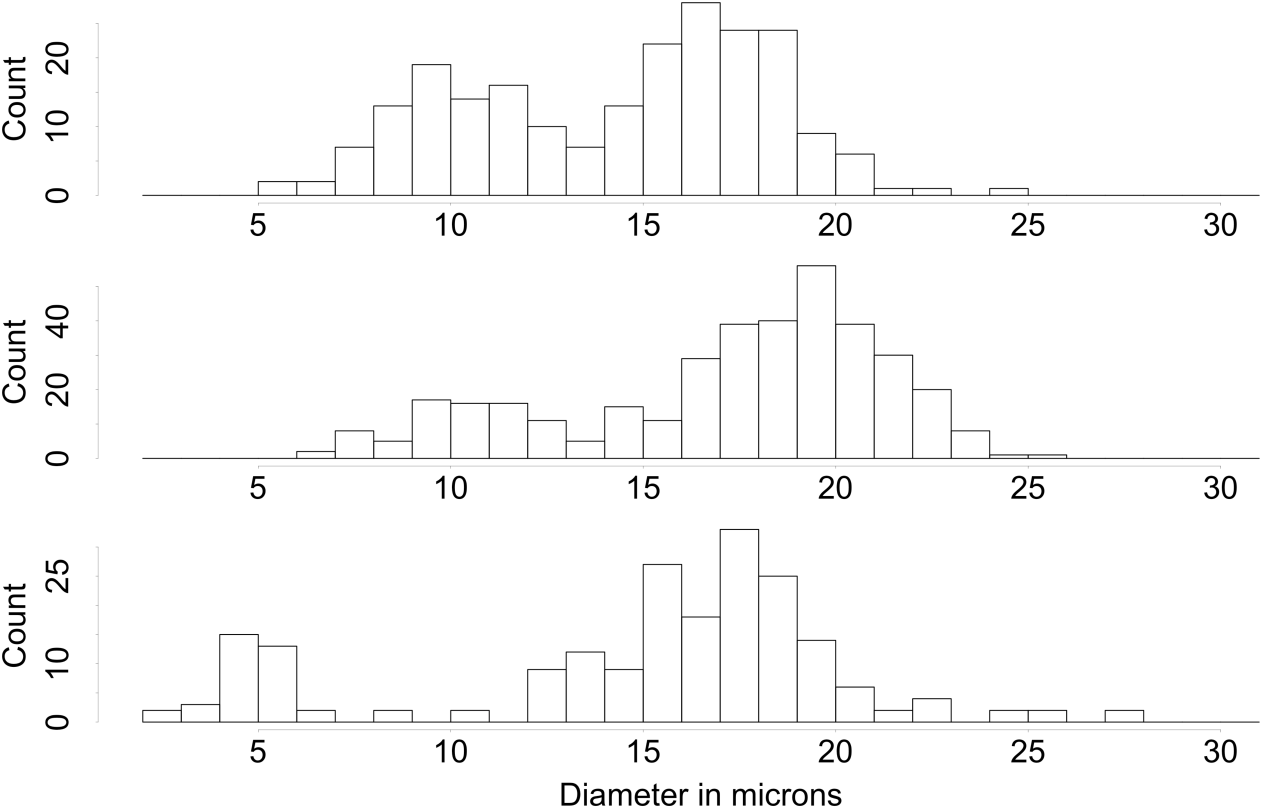
Distribution of aster diameter for *T. authia* samples. From top to bottom: CASIZ 235110, Carpinteria Reef; CASIZ 235111, Perkin’s Reef, Monterey Bay; TLT 364, Saddleback Ridge, Santa Cruz Island.

### Microscopic features

Perpendicular sections show tissue packed with asters (figure 6). Megascleres (styles/tylostyles) are found in projecting bundles, tips up, that pierce the surface of the sponge. Bundles fan-out as they rise through the sponge, resulting in bouquets of spicules at the surface. Other megascleres are found parallel to the substrate in the lower third of the tissue section.

## Geographic distribution

The range of this species in the United States was previously described as Southern California only (Lee *et al*. 2007). This range is here extended North to Monterey Bay in Central California. To the South, samples from as far away as Chile have been assigned to this species, though no molecular data is yet available for confirmation (Carballo and Cruz-Barraza 2006, Desqueyroux-Faúndez 1972). The depth range of the species was previously described as “intertidal to deep water” (Lee *et al*. 2007). All three samples collected in the current work were from the shallow subtidal, 3.5 – 11 m in depth. It was not found in the intertidal, but sampling in the intertidal was very limited. The holotype was collected from the Southern California intertidal in 1930 (de Laubenfels 1932).

## Remarks

Genetic data strongly support membership for this species in the Tethyidae and not the Timeidae. As discussed in the phylogentic results, it is currently difficult to ascertain the differences between *Timea* and *Tethyatimea*, because the species membership of each genus is incompletely ascertained. One proposed distinction has been the presence of size classes of spicules among the *Tethytimea* (Cruz-Barraza *et al*. 2017, Sará 2002). I find that samples of *T. authia* have a bimodal distribution of aster sizes, consistent with this diagnosis. The overlapping size ranges may partially explain why this sponge has not been diagnosed as a *Tethyatimea* in the past.

Family Hemiasterellidae Lendenfeld, 1889

*Galaxia* gen. nov.

## Type species

*Galaxia gaviotensis* sp. nov.

## Diagnosis

Hemiasterellidae with thinly encrusting growth habit. Megascleres include tylostyles, styles, and oxeas. These are found in bundles parallel to the substrate and oriented vertically, piercing the sponge surface. Microscleres are asters.

## Etymology

From the latin galaxia, referring to the milky way. Like a galaxy, the type species of the genus is packed with a diversity of stars.

*Galaxia gaviotensis* sp. nov.

## Material Examined

Holotype, CASIZ 235112, Arroyo Quemado Reef, Santa Barbara County, California, United States of America (34.46775, -120.11905), 15 meters depth, 7/29/19.

## Etymology

The area where this sponge was collected is known as the Gaviota Coast, and is the longest remaining undeveloped part of the Southern California coastline. As a common name, I suggest “Gaviota galaxy sponge”.

## Description

### Macroscopic features

The sponge formed a pale-yellow crust with surface dimensions approximately 10 ⨯ 5 cm. Thickness only 200-650 μm. Field photos (figure 8) show darker spots on the surface; these disappeared in sampled fragments, so were likely oscules or pores that contracted when disturbed.

**Figure 8.**
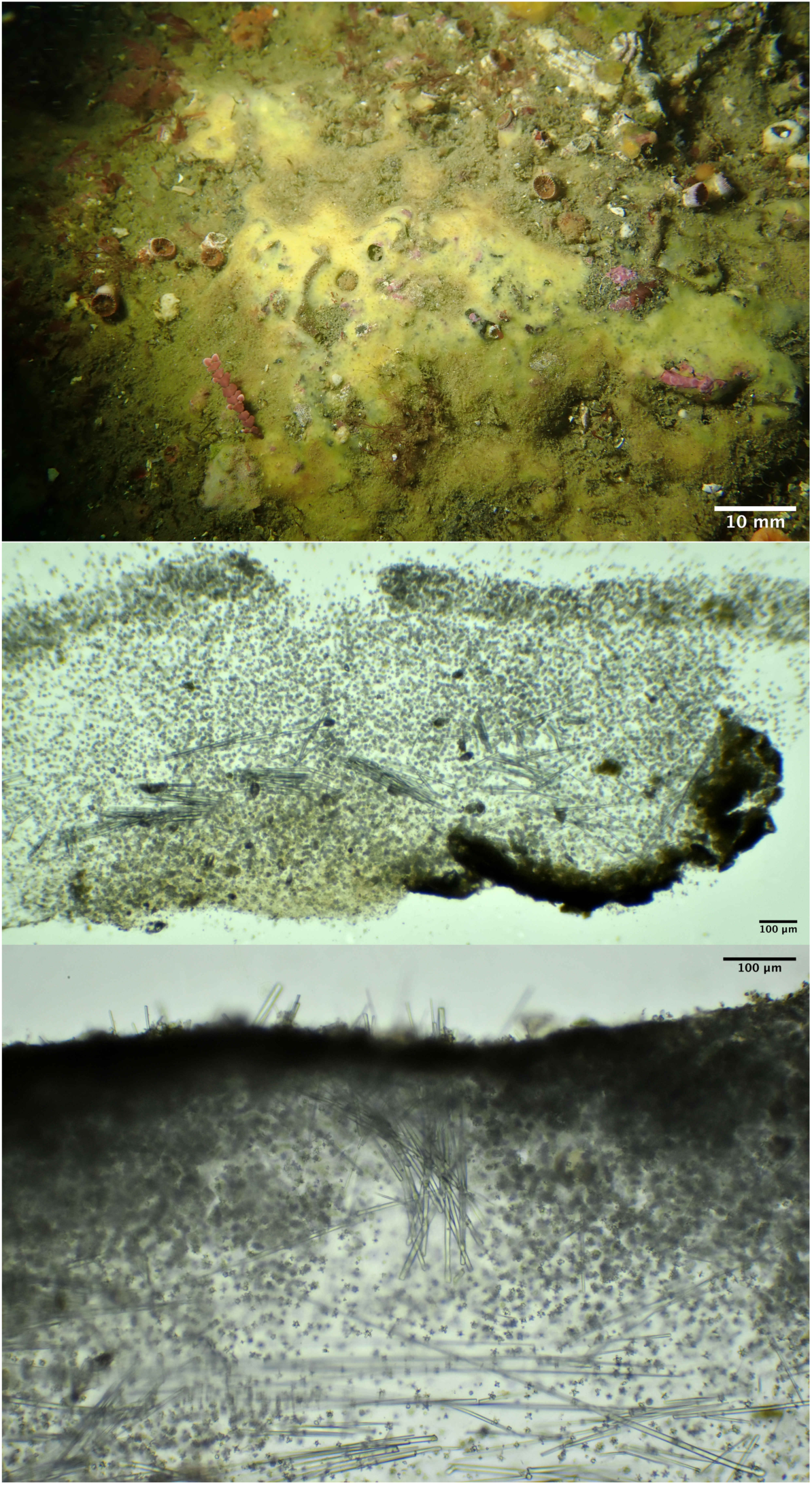
*Galaxia gaviotensis*. Top: sponge in situ before collection. Perpendicular sections (center, bottom) show skeletal morphology of asters and megascleres.

### Microscopic features

Styles comprised 70% of megascleres. Some were tylostyles with well-formed oval heads, while others had weak heads (subtylostyles), and others had lumpy, irregular heads (fig. 9). Many also had subterminal heads or weak bands of slightly increased width near the head of the style. About 13% were unadorned, simple styles. Length varied greatly, from 176-1330 μm (mean = 751, N=45). Width from 2-9 μm, with a mean of 5.3 μm. The other 30% of megascleres were asymmetrical oxeas. These were sharply pointed at both ends, and gently bent at the thickest point, which was closer to one end. A minority (20%) of these had very weak centrylote swelling at the point of inflection. Length was less variable than for styles, from 174-464 μm (mean = 258, N=19); width 2-5 μm, with a mean of 3.7 μm.

**Figure 9.**
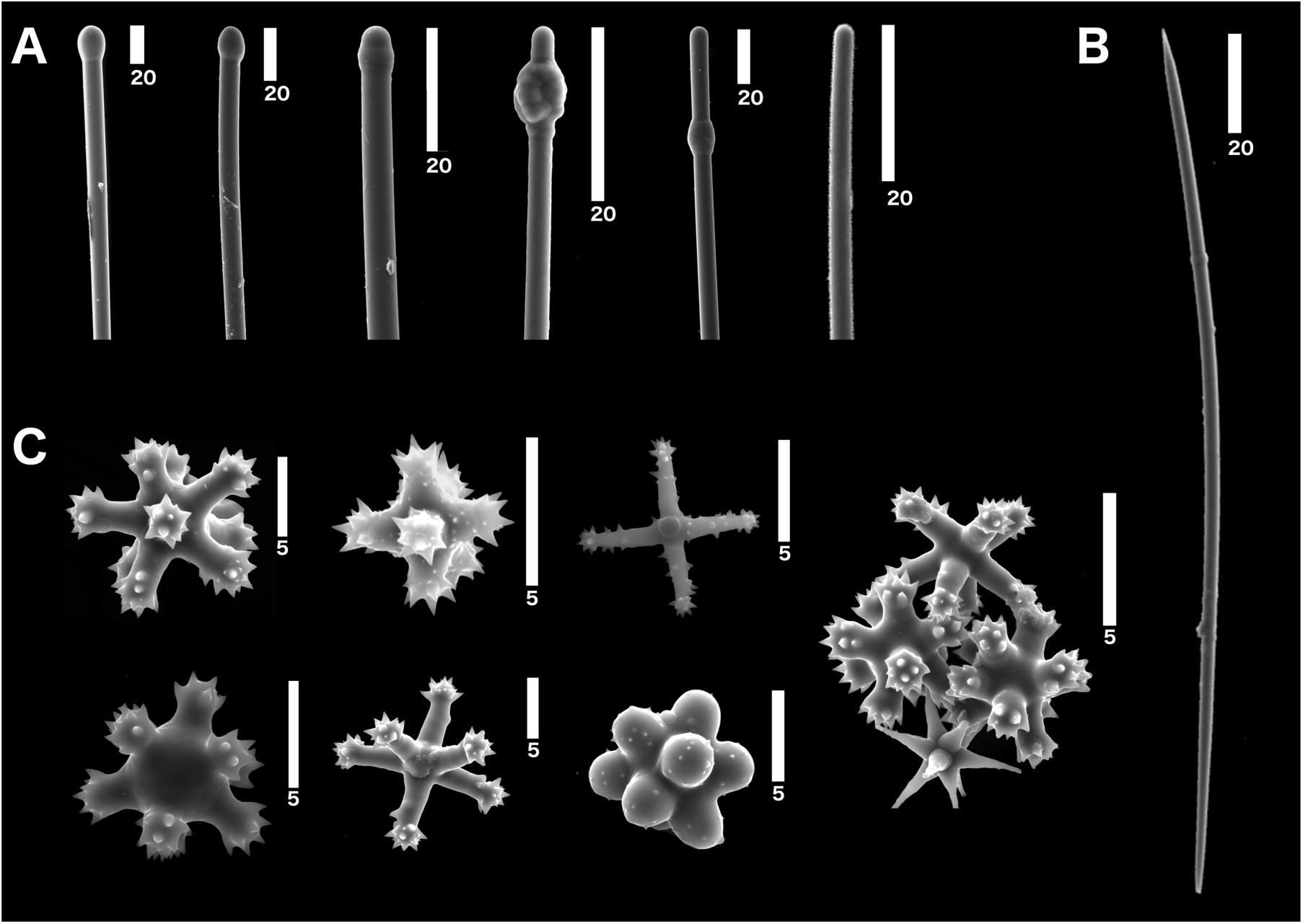
Morphology of *G. gaviotensis* spicules. A) Tylostyles and styles; B) oxea; C) euasters. Numbers below scale bars indicate size in μm.

Microscleres consisted of abundant and varied euasters. The most common type had thick, untapered rays ending in enlarged, spiny bouquets (“tylasters”). The ratio of ray length to centryole size varied greatly (fig. 9). Some were also spined along the rays, instead of merely at the tips. In some asters, the rays were greatly reduced, to hemispherical bulges around a large centrum, sometimes bearing spines and sometimes nearly smooth. Asters with conical rays (oxyspherasters) were also present. These ended in spined bouquets as well, but the bouquets were sometimes very small, resulting in nearly smooth, tapering, conical rays. Mean aster diameter was 10.6 μm (range 4 - 22 μm, N=62); sizes were not clearly different among types.

### Spicule arrangement

Perpendicular sections through the sponge tissue revealed that it was packed with asters (figure 8). These formed a thick surface crust, but were also abundant throughout; no pattern of arrangement by type could be discerned. Megascleres were most dense in the lower half of the sponge, running parallel to the substrate; many were found in bundles. Bundles of megascleres were also oriented vertically, points up, piercing the surface of the sponge.

## Geographic distribution

This sponge has been found at only a single location: Arroyo Quemado reef, approximately 0.15 km offshore of the small community of Arroyo Quemado (34.468801, -120.121233). This is an area of rocky reef and kelp forest that is part of the Santa Barbara Coastal Long-Term Ecological Research Station (sbclter.msi.ucsb.edu). It was found on “transect 3”, which is a flat area 7.5 - 9.5 m deep, strewn with cobbles and boulders. It was growing on a single boulder. I made a total of 8 collecting dives at this reef (5 hours of cumulative search time) but no additional individuals were found. Seven months after collecting the initial sample, the transect 3 area was re-visited in an attempt to relocate the original sponge, without success. Many of the boulders at the site were coated in a fine layer of silt at this time, making it likely that the sponge was still present but concealed.

## Remarks

Most encrusting sponges in the Tethyida are placed in the genus *Timea. Timea* is rejected in the current case due to the presence of oxeas and genetic data that places the species in the Hemiasterellidae. Encrusting Hemiasterellidae are known from only two genera: *Leptosastra* and *Liosina. Leptosastra* is monotypic, with the unusual *L. constellata* having acanthostyles, and tornotes; this is a very poor fit for the current species. *Liosina* includes at least one encrusting species (though thickly encrusting, with chimneys) and sometimes have oxeas; however, these species lack asters. *Hemiasterella* include some species with oxeas, but these species are all massive, digitate, or vase-shaped. Thus, this species has become the type for a new genus.

## Conclusions

By combining molecular and morphological approaches, this work significantly improves our understanding of Californian Tethyida. Taxonomic updates include the discovery and description of *G. gaviotensis* gen nov. sp. nov., the reassignment of *Timea authia* to the Tethyidae as *Tethytimea authia* comb. nov., and the synonymy of *Tethya californiana* and *Tethya vacua*. Genetic data were key to these revisions, and, in my opinion, are indispensable in poriferan taxonomy in general. It seems unwise to erect a new species based solely on the presence of a single morphological character with unproven utility, like the presence of vacuoles. However, it is also unfortunate that morphological data is not publicly available for many of the sequenced vouchers in Genbank. This greatly reduces the utility of publicly available sequence data. Here, I have provided a combination of morphological and genetic data that will be useful for ongoing efforts to revise poriferan taxonomy. My hope is that future work on the ecology and evolution of these species improves both our understanding of the near-shore California ecosystem and the evolution of these interesting animals.

## Acknowledgements

I am grateful for the help and support of many people in UCSB’s Marine Science Institute and Diving & Boating Program, especially Robert Miller, Clint Nelson, Christoph Pierre, Frankie Puetzer, Christian Orsini, H. Mark Page, and Alecia Dezzani. Steve Lonhart (NOAA) and Shannon Myers (UCSC) were instrumental in facilitating collections in Central California, and the Natural History Museum of Los Angeles’ DISCO program facilitated collections in Los Angeles County. Christina Piotrowski (California Academy of Sciences) and Patricia Gómez (National Autonomous University of Mexico) graciously shared vouchers from their collections.

## Funding declaration

Financial support was provided by UCSB and by the National Aeronautics and Space Administration Biodiversity and Ecological Forecasting Program (Grant NNX14AR62A); the Bureau of Ocean Energy Management Environmental Studies Program (BOEM Agreement MC15AC00006); the National Oceanic and Atmospheric Administration in support of the Santa Barbara Channel Marine Biodiversity Observation Network; and the U.S. National Science Foundation in support of the Santa Barbara Coastal Long Term Ecological Research program under Awards OCE-9982105, OCE-0620276, OCE-1232779, OCE-1831937. The funders had no role in study design, data collection and analysis, decision to publish, or preparation of the manuscript.

---

**Supplementary table 1.** Sampling effort and locations. Intertidal/marina sites are listed last. For all subtidal sites, the number of sampling dives is listed, and the cumulative dive time as an estimate of search effort.

| Region | Location name | Lat | Long | Dives | Max depth (m) | Cumulative search time (min) | Tethya californiana | Tethytimea authia | Galexia gaviotae |
| --- | --- | --- | --- | --- | --- | --- | --- | --- | --- |
| Monterey | Middle Reef, Pt. Lobos | 36.52172 | -121.93894 | 1 | 15 | 60 |  |  |  |
| Monterey | Hopkin's Marine Station | 36.62084 | -121.90142 | 1 | 11 | 74 | Collected |  |  |
| Monterey | Perkin's Reef | 36.62920 | -121.92031 | 1 | 12 | 46 | Photo | Collected |  |
| Santa Barbara | Arroyo Hondo Reef | 34.47182 | -120.14262 | 2 | 8 | 52 |  |  |  |
| Santa Barbara | Arroyo Quemado Reef | 34.46775 | -120.11905 | 7 | 11 | 302 | Collected |  | Collected |
| Santa Barbara | Tajigus | 34.46279 | -120.10185 | 2 | 8 | 93 |  |  |  |
| Santa Barbara | Naples Reef | 34.42212 | -119.95154 | 6 | 16 | 279 | Collected |  |  |
| Santa Barbara | Elwood Bluffs | 34.41775 | -119.90150 | 3 | 14 | 138 | Collected |  |  |
| Santa Barbara | Coal Oil Point | 34.40450 | -119.87890 | 5 | 11 | 257 | Collected |  |  |
| Santa Barbara | Isla Vista Reef | 34.40278 | -119.85755 | 4 | 12 | 135 |  |  |  |
| Santa Barbara | Goleta Sewer Pipe | 34.41420 | -119.82930 | 1 | 6 | 22 |  |  |  |
| Santa Barbara | Mohawk Reef | 34.39407 | -119.72957 | 7 | 9 | 220 | Collected |  |  |
| Santa Barbara | 1000 Steps | 34.39472 | -119.71347 | 1 | 7 | 30 |  |  |  |
| Santa Barbara | Leadbetter Reef | 34.39895 | -119.69703 | 1 | 9 | 14 |  |  |  |
| Santa Barbara | Oil Platform C | 34.33293 | -119.63173 | 1 | 23 | 39 |  |  |  |
| Santa Barbara | Oil Platform A | 34.33189 | -119.61353 | 1 | 27 | 35 |  |  |  |
| Santa Barbara | Carpinteria Reef | 34.39163 | -119.54169 | 2 | 6 | 61 |  | Collected |  |
| Santa Barbara | Elwood Junkpile | 34.42687 | -119.92390 | 1 | 12 | 34 |  |  |  |
| Santa Barbara | UCSB intake pipe | 34.40297 | -119.84080 | 1 | 18 | 34 |  |  |  |
| Los Angeles | Hawthorne reef | 33.74714 | -118.42090 | 1 | 24 | 39 | Photo |  |  |
| Los Angeles | Halfway reef | 33.76265 | -118.42560 | 1 | 23 | 40 |  |  |  |
| Los Angeles | Resort Wall | 33.76499 | -118.42815 | 1 | 21 | 43 |  |  |  |
| Los Angeles | Flat rock | 33.79662 | -118.41110 | 1 | 8 | 45 |  |  |  |
| Los Angeles | Redondo barge | 33.83833 | -118.41040 | 2 | 27 | 49 |  |  |  |
| San Diego | Goalpost | 32.69438 | -117.26860 | 1 | 15 | 50 | Photo |  |  |
| San Diego | Six Fathoms | 32.71000 | -117.26860 | 1 | 18 | 40 | Photo | Photo |  |
| San Miguel Island | Wreck of the Cuba | 34.03009 | -120.45575 | 1 | 10 | 52 | Photo |  |  |
| San Miguel Island | Inside of Wycoff Ledge | 34.02132 | -120.38710 | 1 | 19 | 52 | Photo |  |  |
| San Miguel Island | Miracle Mile, E of Crook Point | 34.01367 | -120.34180 | 2 | 20 | 108 | Photo |  |  |
| Santa Rosa Island | Johnson's Lee | 33.89966 | -120.10735 | 2 | 15 | 65 | Collected |  |  |
| Santa Cruz Island | Hazzard's, Santa Cruz Is. | 34.06066 | -119.82810 | 1 | 13 | 75 |  |  |  |
| Santa Cruz Island | Cueva Valdez, Santa Cruz Is. | 34.05498 | -119.81960 | 1 | 14 | 75 |  |  |  |
| Santa Cruz Island | Platt's Harbor, Santa Cruz Is. | 34.04765 | -119.73537 | 1 | 10 | 86 |  |  |  |
| Santa Cruz Island | Marmeda, Santa Cruz Is. | 33.98378 | -119.6391 | 1 | 13 | 35 | Collected |  |  |
| Santa Cruz Island | E. of Marmeda, Santa Cruz Is. | 33.98380 | -119.61290 | 1 | 15 | 38 |  |  |  |
| Santa Cruz Island | The Suburbs, Santa Cruz Isl. | 34.05291 | -119.58309 | 1 | 14 | 59 |  |  |  |
| Santa Cruz Island | Big Rock, Santa Cruz Is. | 34.05220 | -119.57360 | 5 | 14 | 299 | Photo |  |  |
| Santa Cruz Island | Pink Ribbon | 33.98967 | -119.52470 | 1 | 11 | 60 |  |  |  |
| Santa Cruz Island | Saddleback Ridge | 34.03817 | -119.52470 | 2 | 12 | 134 |  | Collected |  |
| Anacapa Island | West End, Anacapa Is. | 34.01352 | -119.44570 | 1 | 12 | 40 | Collected |  |  |
| Anacapa Island | Zebra Cove | 34.01000 | -119.44000 | 1 | 13 | 66 | Photo |  |  |
| Anacapa Island | Landslide Cove, Anacapa Is. | 34.01539 | -119.43560 | 1 | 11 | 49 |  |  |  |
| Anacapa Island | Underwater Island, Anacapa Is. | 34.00720 | -119.42959 | 2 | 18 | 102 |  | Photo |  |
| Anacapa Island | Cat Rock | 34.00426 | -119.42335 | 1 | 11 | 65 |  |  |  |
| Anacapa Island | Portuguese Rock, Anacapa Is. | 34.01545 | -119.42149 | 2 | 26 | 111 |  |  |  |
| Anacapa Island | Underwater Arch, Anacapa Is. | 34.01399 | -119.36224 | 1 | 10 | 45 |  |  |  |
| Anacapa Island | Landing Cove, Anacapa Is. | 34.01690 | -119.36090 | 1 | 15 | 57 |  |  |  |
| Catalina Island | Parson's Landing | 33.47502 | -118.55000 | 2 | 17 | 120 | Photo |  |  |
| Catalina Island | Eel Cove | 33.45662 | -118.51120 | 1 | 9 | 30 |  |  |  |
| Monterey | Point Pinos Intertidal | 36.63447 | -121.93923 | n/a | n/a | 90 |  |  |  |
| Monterey | Monterey Marina | 36.60797 | -121.89288 | n/a | n/a | 20 |  |  |  |
| Santa Barbara | COP Intertidal | 34.406656 | -119.8775 | n/a | n/a | 60 |  |  |  |
| Santa Barbara | Santa Barbara Harbor | 34.405592 | -119.68964 | n/a | n/a | 150 |  |  |  |
| Los Angeles | 22nd Street Marina | 33.724772 | -118.28078 | n/a | n/a | 20 |  |  |  |
| San Diego | Point La Jolla | 32.85112 | -117.27298 | n/a | n/a | 60 |  |  |  |
| San Diego | Mission Bay Harbor | 32.764209 | -117.23829 | n/a | n/a | 40 |  |  |  |

**Supplementary table 2.**
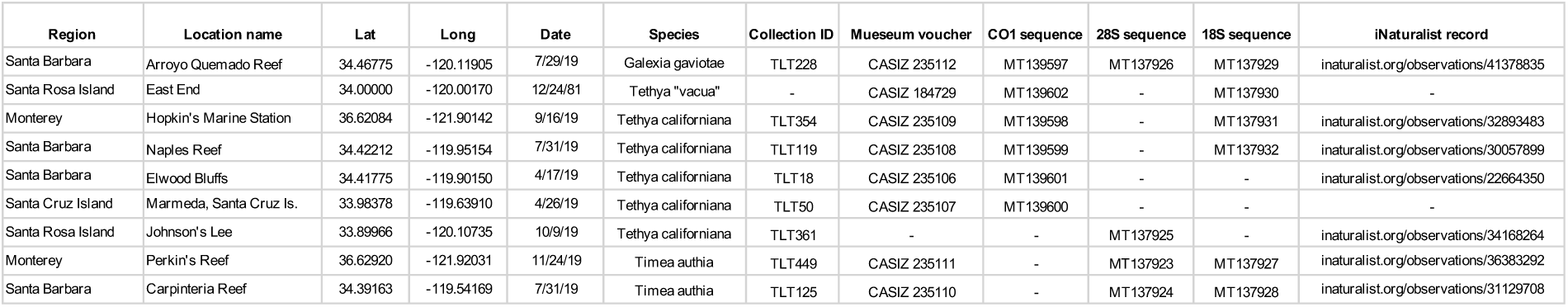
Archival information for vouchered samples and samples with sequence data. CASIZ: California Academy of Sciences Invertebrate Zoology.

Supplementary table 3. Raw data on spicule dimensions for all samples.

(This data set is not easily formatted to fit into a text document but is available upon request, and will be submitted to the journal upon acceptance of the manuscript.)

Supplementary table 4. Genetic metadata.

(This data set is not easily formatted to fit into a text document but is available upon request, and will be submitted to the journal upon acceptance of the manuscript.)

## Notes

### Competing Interest Statement

The authors have declared no competing interest.

